# Performance in a GO/NOGO perceptual task reflects a balance between impulsive and instrumental components of behaviour

**DOI:** 10.1101/051607

**Authors:** A. Berditchevskaia, R.D. Cazé, S.R. Schultz

## Abstract

In recent years, simple GO/NOGO behavioural tasks have become popular due to the relative ease with which they can be combined with technologies such as *in vivo* multiphoton imaging. To date, it has been assumed that behavioural performance can be captured by the average performance across a session, however this neglects the effect of motivation on behaviour within individual sessions. We investigated the effect of motivation on mice performing a GO/NOGO visual discrimination task. Performance within a session tended to follow a stereotypical trajectory on a Receiver Operating Characteristic (ROC) chart, beginning with an over-motivated state with many false positives, and transitioning through a more or less optimal regime to end with a low hit rate after satiation. Our observations are reproduced by a new model, the Motivated Actor-Critic, introduced here. Our results suggest that standard measures of discriminability, obtained by averaging across a session, may significantly underestimate behavioural performance.

---

What is the impact of motivation on behaviour? Reinforcement learning theory assumes that an animal optimises behaviour according to the value placed on the goal of an action under different levels of deprivation (Dayan and Niv 2008; Guitart-Masip et al. 2010). Beyond the value placed on the goal (*directional effect*), motivational state is also critically important in regulating the overall effort and rate of activity (*activating effect)* an animal engages in (Niv et al. 2007; Salamone and Correa 2012). The combination of behavioural training with sophisticated electrophysiological and imaging techniques is beginning to provide unprecedented insight into the functioning of neural circuits underpinning perceptually-driven decision making (Huberman and Niell 2011; Carandini and Churchland 2013). A commonly employed paradigm for studying perceptual decisions is the two-category GO/NOGO task, where the animal performs a response to obtain a reward during a ‘Go’ stimulus, and needs to withhold the response for the ‘NoGo’ stimulus. This paradigm has been commonly used in recent years to draw inferences about sensory computations (Lee et al. 2012; Glickfeld et al. 2013; Pinto et al. 2013; Bracey et al. 2013; Fu et al. 2014; Zhang et al. 2014). Mice are readily able to perform such tasks, however natural biases (Busse et al. 2011) can interfere with performance. Motivational influences on behaviour have also been documented both in rodents (Niv 2007, Komiyama et al. 2010) and other species (Mayack and Naug 2015). These factors increase the difficulty of devising appropriate training protocols and can impact on the interpretation of results (Carandini and Churchland 2013).

The typical timeline for observation during simple decision-making tasks in modern neuroscience extends to hundreds of trials, thus capturing a range of motivational levels throughout a single behaviour session. While there have been mentions of changes in motivation within individual sessions previously (Andermann et al. 2010, Stüttgen et al. 2011), these effects are often ignored (Niv et al. 2007; Busse et al. 2011) or factored out of analyses (Rivalan et al. 2013). Here, we have analysed the effect of motivation on a GO/NOGO visual discrimination task at the single session level. To our knowledge, this is the first detailed exploration of these factors within the context of the GO/NOGO paradigm, although previously published work (Komiyama et al. 2010) can be re-examined to support our conclusions. Our approach enables us to evaluate the role of motivation on classification performance, which has important implications for perceptual computation and decision-making.

Our results demonstrate that motivational state changes throughout the course of a session in a quantifiable and consistent manner, as measured by response vigour (lick frequency). We show that an overly high level of motivation interferes with behaviour, causing poor performance until initial thirst is satisfied. Lick latency analysis allows further decomposition of the response into different behavioural components, aiding understanding of performance on the task. We conclude that while learning is likely to be reliant on maintenance of water restriction, it is more effective to keep animals in a more satiated state thereafter to gain accurate insights into behaviour.

## Results

### Overmotivation masks true sensitivity at the beginning of a session

We trained mice (n=15) under water restriction (see Methods) to perform a visual discrimination task, using vertical (S+) and horizontal (S-) square-wave gratings initially projected to the entire visual field (full-field) and then the monocular region of the visual field (Figure 1). After reaching criterion and showing a high level of discrimination during 3 consecutive sessions of the monocular behavioural task, we recorded the performance of the mice during a test session. Importantly, at this stage all animals were at a comparable level of performance. Figure 2a shows a typical example of a lick report from a post-criterion behavioural session. We define a single behaviour session as the 30-60 minute period where mice actively participated in the discrimination task. Despite slight differences in the absolute values of Hit and False Alarm (FA) rates, all mice showed the same pattern of changes to their licking behaviour over the course of a session. To analyse these changes, we split each session into three equal segments (Initial, Middle, Final). Our aim was to examine the impact of motivation on the expression of a learned instrumental behaviour on the timescale of a single session (within-session changes). We normalised the length of the three segments to the total number of trials, for each mouse, and used standard signal detection theory (SDT) measures of Hit and FA rates in each segment (see Methods).

**Figure 1.**
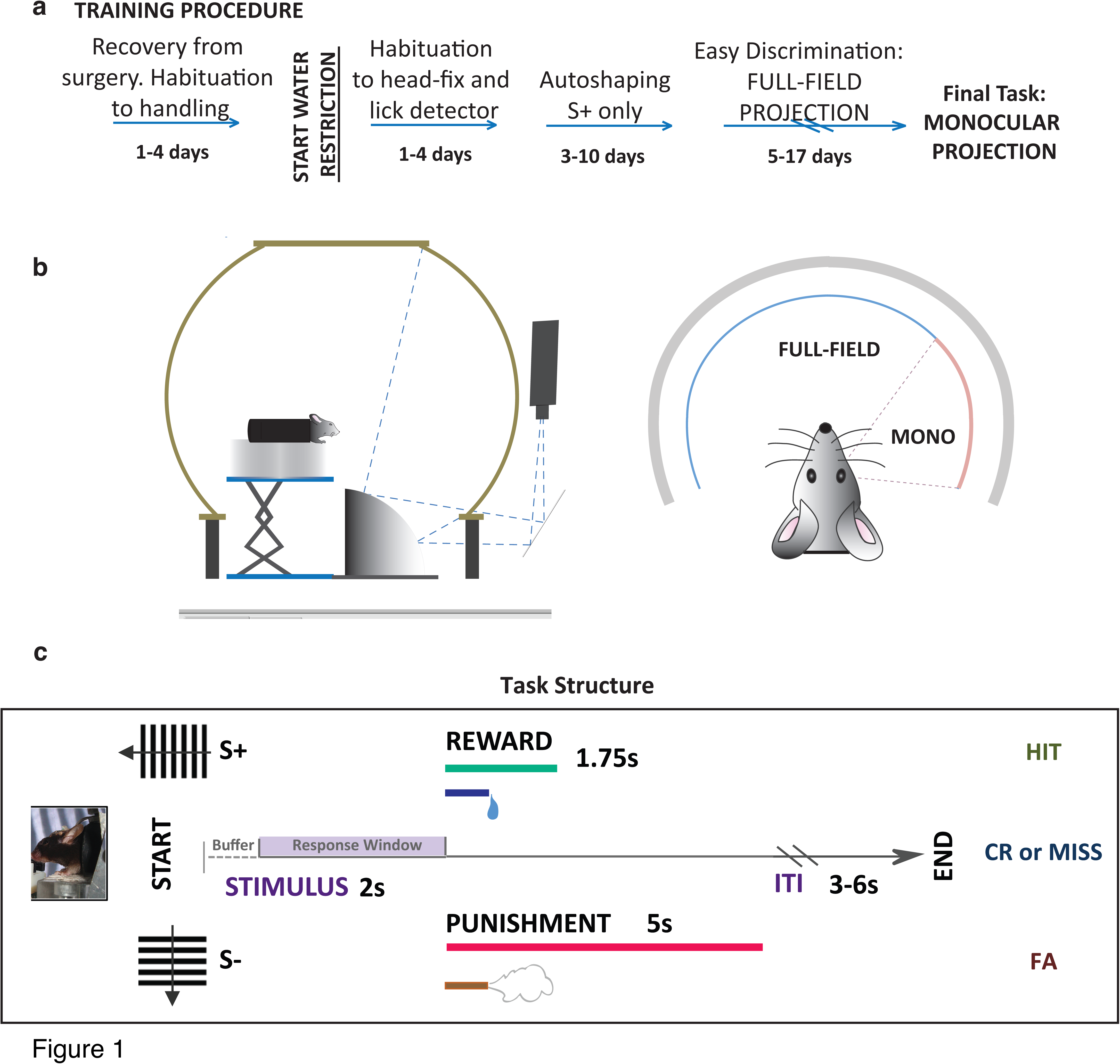
Go/NoGo visual discrimination task and training procedure. ***a***. Full behavioural training and experimental timeline. ***b***. All experiments were carried out under head-fixed conditions in an immersive dome environment. During initial training, stimuli were projected onto the entire inner surface of the dome (FULL-FIELD), before restriction to a monocular (MONO) projection region for the final task. ***c***. Task sequence showing all possible trial and outcome types with timing. Small rectangles (ie, orthogonal gratings, black, grey) show image projected onto screen during each trial state. A lick during the Stimulus (up to 2s) state triggered immediate transition into a Reward (1.75s) or Punishment (5s) state, followed by a variable length (3-6s) ITI state. *S+ = Go stimulus, positively reinforced with water; S-= NoGo stimulus, negatively reinforced with airpuff; ITI = inter trial interval; buffer = 0.5s at start of trial, licks during this period do not trigger a state change to Reward/Punishment state*.

**Figure 2.**
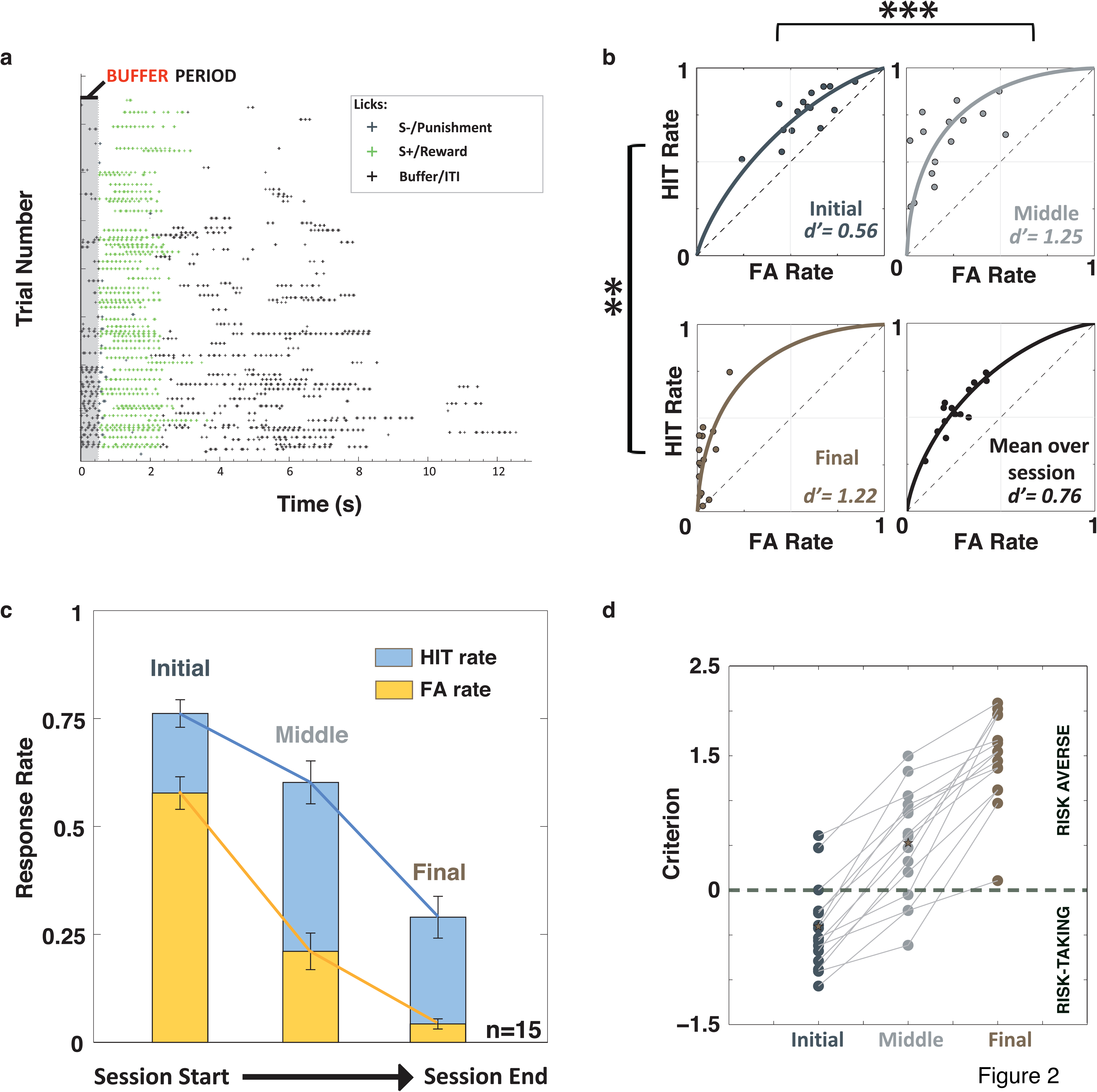
Discrimination sensitivity starts well below its final value, gradually improving as the level of deprivation decreases over the course of a session. ***a***. The behavioural readout from a typical session once criterion had been reached on the final version of the task. Time increases from bottom to top for the session and left to right for single trials. Licks per trial are tracked horizontally on each line, with each coloured cross representing a lick. Vertical orange lines provide a visual reference for subsequent lick latency (*LLat*) results. ***b***. Separate panels show group isosensitivity curves for the Initial, Middle and Final stages of the session. For each section of the session, we calculated the d′ for each of the Hit/FA pairs within that section, and used the mean of these values to generate the curve. The bottom right panel shows the d′ obtained from averaging over the whole session (n=15, data represent mean +/- s.e.m. ** p<0.005, *** p<0.0003, Student’s t-test). ***c***. Hit and FA rates over different stages of the testing session on the final task (n=15).

At the start of the session mice performed with high values for both Hit and FA rates (Figure 2c: Hit_Initial_ = 0.76 ± 0.03, FA_Initial_ = 0.58( ± 0.04). In the Middle segment, Hit rates remained high while FA rates dropped (Figure 2c: Hit_Middle_ = 0.60 ± 0.05, FA_Middle_ = 0.21 ± 0.04). During the Final segment, both response rates decreased (Figure 2c: Hit_Final_ = 0.29 ±0.05, FA_Final_ = 0.04 ± 0.01. In other words, we observed that the Hit and FA rates of water-restricted mice showed dramatic variation over the course of a session, with corresponding changes in the SDT discriminability measure, *d*′. In Figure 2, we show these changes in d′ during the Initial, Middle and Final segments as performance curves in Receiver Operating Characteristic (ROC) space (see Methods). The sensitivity of the mice in the early part of the session was less than half of the value seen in the later segments of the session (*d*′_Initial_ = 0.56 ± 0.07, *d*′_Middle_ = 1.25 ± 0.14, *d*′_Final_ = 1.22 ± 0.16). A comparison of the group values for each of the 3 segments showed that the sensitivity of the mice during the Initial segment was significantly different to the d′ values that emerged in the later stages of the session (Student’s t-test, p<0.0003). The Final and Initial segment d′ distributions were also significantly different (Student’s t-test, p<0.005) but no difference was found between sensitivity values for the Middle and Final segments. This highlights that, despite differences in overall participation levels through the Middle and Final stages of the session, mice performed with consistent sensitivity. The bottom right panel of Figure 2b (purple) demonstrates that mean *d*′ across the session, the most commonly used behavioural sensitivity measure (Andermann et al. 2010; Lee et al. 2012; Pinto et al. 2013), substantially underestimates the true discrimination ability of the mice, with a *d*′ of 0.76 ± 0.04 for the group. The use of means to assess both the within-session performance of individuals and group progression of learning can distort individual acquisition trends and can cause inaccurate estimates of performance (Gallistel et al. 2004).

We conclude from this that stable discrimination sensitivity only emerges after the initial part of a behaviour session in water-restricted mice. The performance of the mice during this initial segment may skew the estimate of discrimination ability when mean discriminability is taken over the whole session. We hypothesise that the changes we observed in mouse behavioural performance were due to fluctuating motivational state. In the remainder of this paper we examine this hypothesis in more detail.

### Behavioural performance can be explained by a motivation-augmented Actor-Critic model

We first used the SDT response bias measure, criterion value, to assess motivation. In a learnt task modelled using SDT, the criterion designates the threshold at which an animal places the decision to “Go”. The negative or positive signage of the criterion reflects the direction of the bias away from a neutral baseline where the animal is equally likely to “Go” or not. Criterion value can thus be understood as reflecting the “directional” aspects of motivation. Figure 2d shows that the criterion value steadily increased over the course of the behaviour session (C_initial_ = −0.42 (±0.12), C_middle_ = 0.51 (±0.16), C_final_= 1.46 (±0.13)), changing from a low criterion (risk-taking) approach to a higher and more conservative (risk-averse) decision policy.

We next explored a modelling approach to test whether a “motivation” variable could account for the stereotyped behaviour we saw during the task. Actor-Critic models are commonly used to describe reinforcement learning in simple two-alternative and operant behavioural tasks (Dayan and Balleine 2002; Dayan and Niv 2008). We compared the performance predicted by two models. The first (AC) was based on the classic Actor-Critic approach with Q-learning (see Methods for details). The second model (MAC) shown in Figure 3a maintains the classic Q-learning framework and initialisation parameters, but also takes motivational state into account. Motivational state is incorporated through a “thirst” variable, which adjusts the Q-values (see Methods) of the Actor-Critic over the course of the behaviour session. The top panels in Figure 3a show the proportion of Hits/FAs and ROC performance predicted by 15 iterations of the models. They predict the performance trajectory that would be seen in animals that have been trained on a Go/NoGo task. We used the baseline assumption that mice had prior knowledge of the task contingencies during the test session, so that S+ stimuli were valued more highly than the S-stimuli (ie, that mice had attained criterion performance) and initialised the starting Q-values accordingly. We estimated the goodness of the predictions made by the models by calculating the Euclidean distance between the experimental data and the simulated results. A perfect match gives a value of 0, while the furthest possible separation gives a value of 2. The performance predicted by the MAC model (distance = 0.15) more closely reflects the behaviour recorded during the experiments than the classic Q-learner (distance = 0.78). Inclusion of a motivation variable reproduced the lower d′ observed during the Initial segment of the session, and gradually changed towards a consistent discrimination level in the Middle and Final segments of the session. The model was also able to capture the change in task participation whilst maintaining discrimination sensitivity for the Middle and Final segments, as seen in the experimental data.

**Figure 3.**
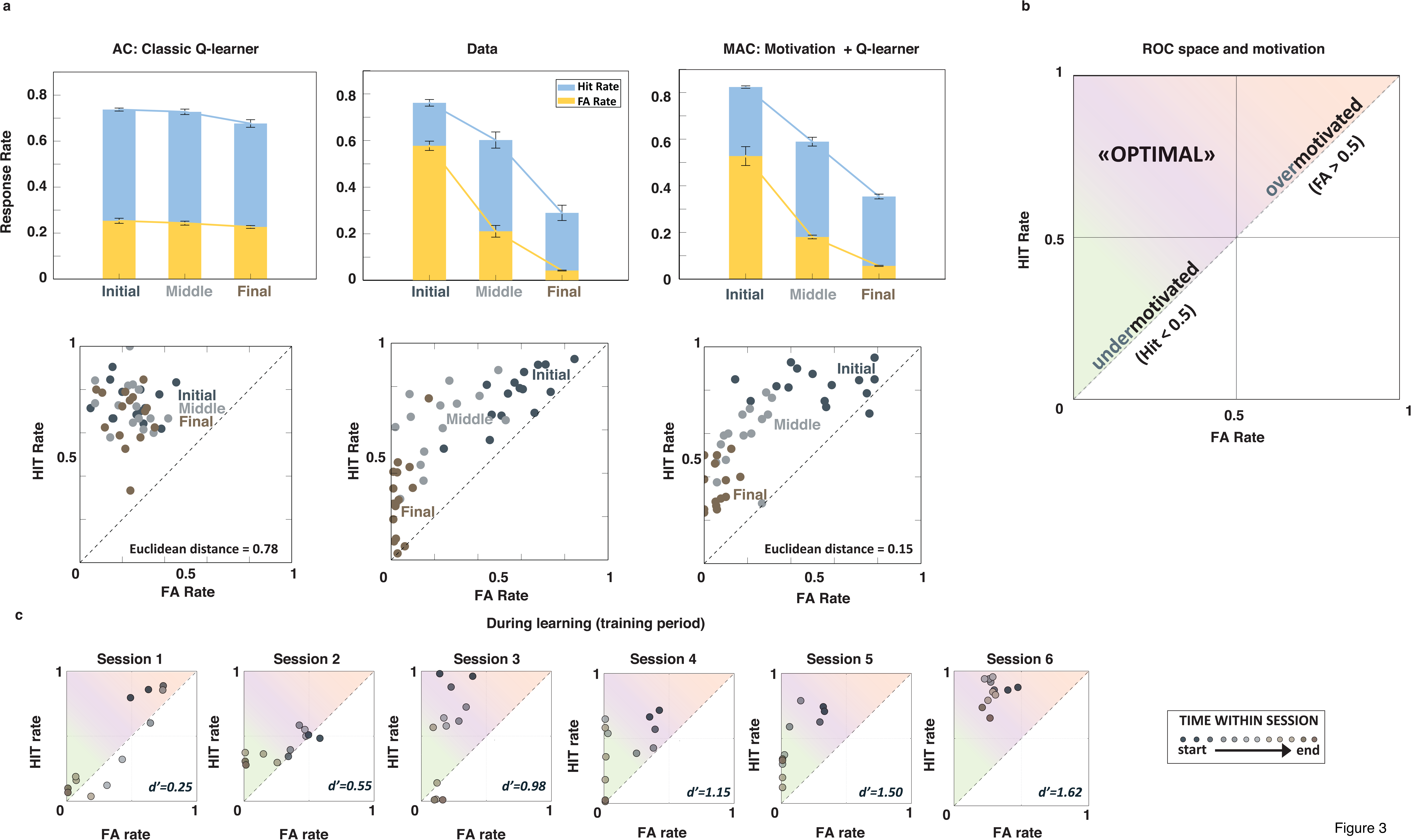
Introducing a motivation variable into the Actor-Critic (AC) model can account for our observations. ***a***. Introducing a motivational factor into the classic Q-learner framework results in performance predictions similar to the changes we observed in the experimental results (Far right panel). Top: while the AC model showed relatively stable sensitivity throughout the entire session, the MAC model recreated the characteristic differences between HIT and FA rates. Bottom: HIT/FA pairs projected into ROC space demonstrate tight clustering for the classic model but spread similar to the experimental results for the motivation model. Modeled data, n=15 iterations, initialized to Q-values of 1 under the assumption that task contingencies had been learnt (see Methods for details). MAC performed better than AC as measured by the Euclidean distance between the values predicted by the model and the experimental results (AC = 0.78; MAC = 0.15). ***b***. Illustrative ROC space: red region shows overmotivated performance, green undermotivated, and purple freedom from extremes of motivational bias. ***c***. The performance of a typical mouse (experimental data) over 6 successive sessions during a training period with full-field stimulation. Each datapoint corresponds to the Hit/FA pair from a given time-point within the session, projected into ROC space. The points move away from the *d′=0* diagonal with learning. The session trajectory varies considerably between sessions, but Hit/FA pairs rarely fall in the overmotivated (red) region of ROC space.

We devised a ROC space classification scheme to assign motivational state (Figure 3b), guided by the ROC literature (Fawcett 2006). Accordingly, when we examined the Hit/FA pairs within ROC space from the experimental test session (Figure 3a: Far Right, bottom panel), we found that behaviour early in the session mostly fell within the section reflecting “high or over-motivation” (red shading) while the Hit/FA pairs from the later stages of the session fell within the purple and green shading regions (“unbiased” and “low motivation”, respectively).

An alternative explanation might posit that mice spend the Initial segment of the session re-learning the task contingencies, until they are able to perform again at a high standard. We compared the typical motivational variation shown by mice during the learning stage of the behavioural training to get a better understanding of the interaction of these two processes. Figure 3c shows 6 consecutive training sessions over which the d′ improved to fulfil the criterion level of performance (First session d′=0.25, Final session d′=1.62). Unlike the learnt stage of behaviour where performance followed a typical trajectory, the performance during the learning phase showed substantial variation in which portion of motivation ROC space the Hit/FA pairs occupied over the sessions. Hit/FA rate values during learning were rarely found in the extreme upper right of the ROC space (red shading). Thus the deprivation regime did not result in overmotivated performance during the learning phase of behavioural training.

Our modelling results thus strongly support a causal role for motivational state in determining the performance trajectory observed on a GO/NOGO task within individual behaviour sessions. In the next section we seek to define these changes in a more quantitative way by considering the *activating* aspects of motivation.

### Quantitative measures of response vigour reliably demonstrate motivational changes within individual sessions

Response vigour, or the rate at which instrumental actions are undertaken, has recently been used as a readout of motivational state (Niv et al. 2007; Salamone and Correa 2012). We quantified response vigour using three complementary measures: lick frequency, lick ‘efficiency’ (or relative licking) and lick latency (Table 1) to characterise changes to motivational state. Licks occurring at specific, well-defined points within a trial can be indicative of different motivational factors and/or changes to behavioural policy when examined in relation to overall lick rates.

**Table 1.**
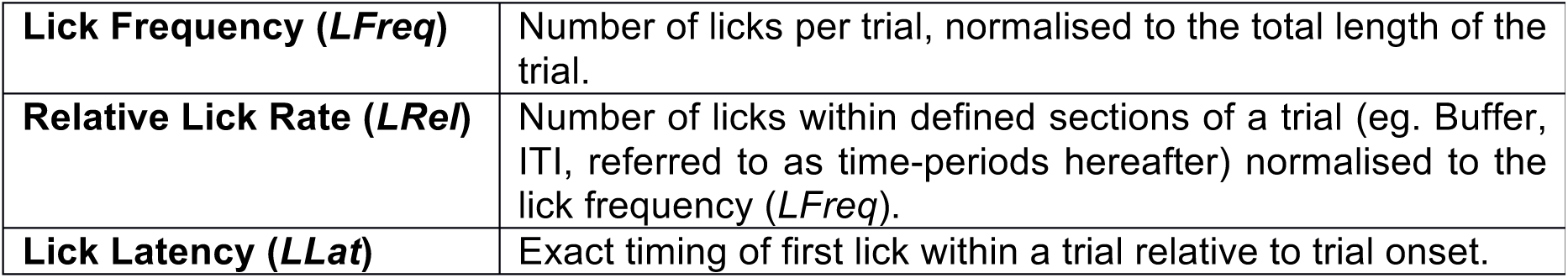
Quantitative measures of response vigour used to measure changes in motivation.

|  |  |
| --- | --- |
| **Lick Frequency (*LFreq*)** | Number of licks per trial, normalised to the total length of the trial. |
| **Relative Lick Rate (*LRel*)** | Number of licks within defined sections of a trial (eg. Buffer, ITI, referred to as time-periods hereafter) normalised to the lick frequency (*LFreq*). |
| **Lick Latency (*LLat*)** | Exact timing of first lick within a trial relative to trial onset. |

We measured the licking frequency (*LFreq*) of the mice during the criterion sessions by counting the absolute number of licks that each mouse performed per trail normalised to the trial length. We then calculated the mean *LFreq* for the ‘Initial’, ‘Middle’ and ‘Final’ segments for both individual mice and the group (Figure 4a). The variability of absolute *LFreq* values within the group was large for the ‘Initial’ segment. We found no systematic trend between the size of these values and deprivation parameters (results not shown) suggesting that this variation was more indicative of an intrinsic baseline of individual animals. Despite this variability all mice followed a group trend, with *LFreq* decreasing over the course of the session (*LFreq*_Initial_ = 4.50 ±0.46, *LFreq*_Middle_ = 3.22 ±0.42, *LFreq*_Final_ = 1.01 ±0.10). This is consistent with the motivational state making a transition between over and undermotivation, where under high motivation animals invigorate an instrumental action (activating effect) that leads to a desirable outcome (directional effect) (Salamone and Correa 2012).

**Figure 4.**
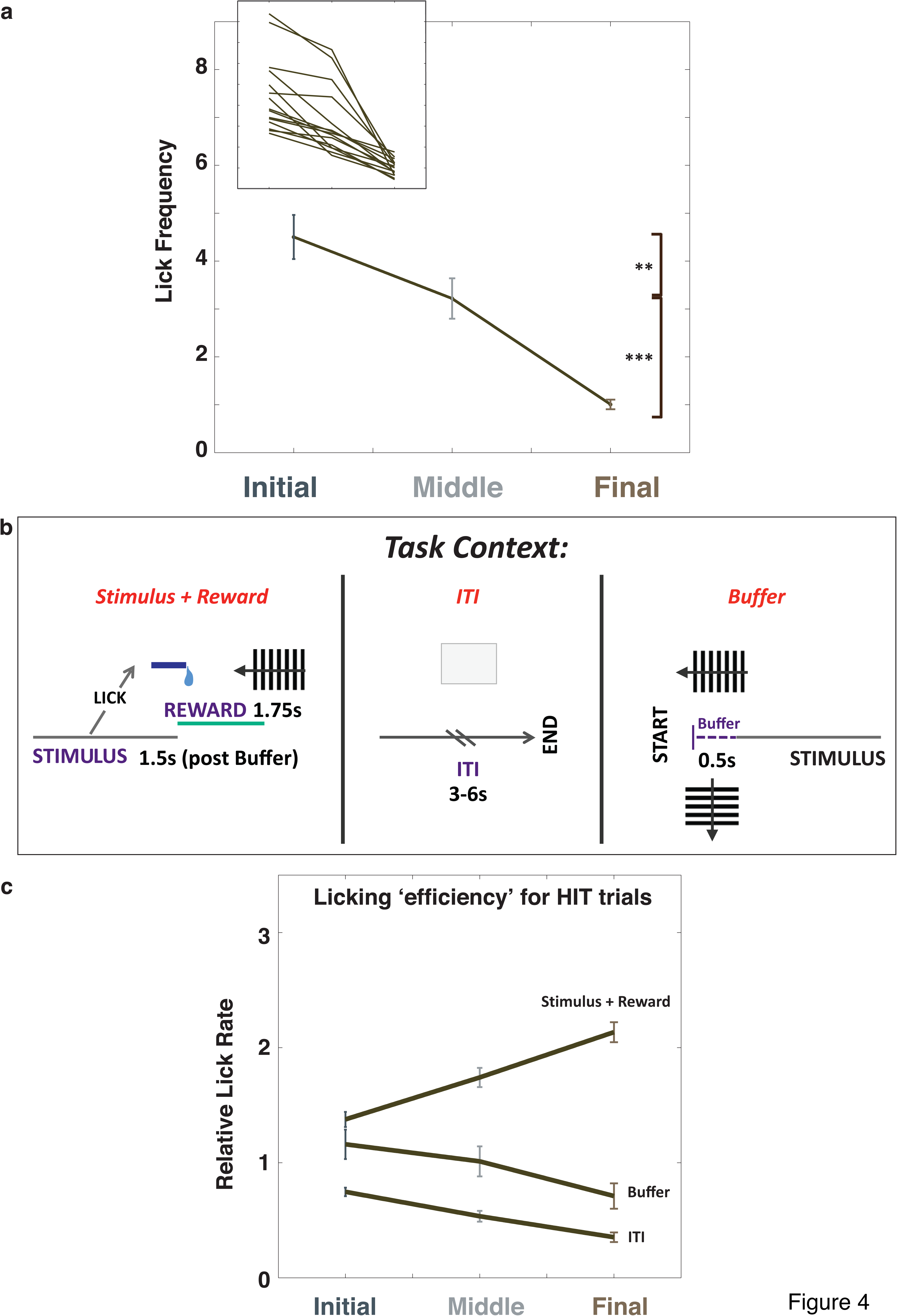
Variation of the lick frequency depends on session progression and task context. ***a***. The rate of the licking action *(Lfreq)* per trial decreases over the session. (n=15, data represent mean +/- s.e.m. ** p<0.0056, *** p<1.05X10-^9^, repeated measure ANOVA with post-hoc comparisons). Inset: *LFreq* for individual mice. Background shading on the main panel corresponds to the 3 different segments of the session (dark grey=Initial, light grey=Middle, brown=Final). ***b***. The lick rate for different periods of a trial normalised to the overall *LFreq* (ie., *LRel*=Relative lick rate). Upper panel shows a schematic of the periods within the trial over which the *LRel* values were calculated: Hits, ITI, Buffer. All *LRel* values were calculated using the trials with a “Hit” outcome. ***c***. *LRel* for Hit, Buffer and ITI time periods over the course of a session. *LRel* for the Buffer and Stimulus+Reward periods are not significantly different during the Initial segment (p=0.75, repeated measure ANOVA with post-hoc comparisons).

We also calculated the relative lick rate (*LRel*) during different time-periods, i.e. the number of licks occurring during a particular time-period divided by the total number of licks in the trial. By this definition, a highly efficient animal would concentrate licking activity into time-periods when the action could lead to triggering or collecting the water and not engage in adjunctive licking in other time-periods. Highly motivated but inefficient animals, on the other hand, may show a more general increase in instrumental response (Dickinson and Balleine 1994) by increasing licking indiscriminately in all time-periods.

For each stage, we compared how *LRel* changed over the session for rewarding time-periods versus those that carried no instrumental gain (Stimulus+Reward vs Buffer or inter-trial interval (ITI) for “Hit” trials, Figure 4b). Figure 4c shows that at the start of the session, the Buffer period (Initial: 1.16 ± 0.13) had similar *LRel* values to the Stimulus+Reward (Initial: 1.38 ± 0.07) period, which were not significantly different (p=0.75). Over the course of the session, these two values separated and the animals licked much more efficiently. This is shown in the significantly higher *LRel* values for the Stimulus+Reward condition than the time-periods where licking held no behavioural advantage (*LRel*_Final_), indicating that animals limited their licking to the instrumentally rewarding periods of the trial.

Comparisons with the time-periods where licking did not lead to a water reinforcement highlighted an interesting difference between the two conditions. We noticed that while *LRel* values during the “hits” time-periods gradually increased, *LRel* in the Buffer period decreased over the course of the session (Initial: 1.16 ± 0.13, Middle:1.01 ± 0.13, Final:0.62 ± 0.11). In contrast, the *LRel* values for the ITI period showed more stability over the session and less variability between mice (Initial: 0.75(±0.04), Middle: 0.53(±0.05), Final: 0.35(±0.04)), suggesting they represented a baseline licking tendency while Buffer licking was more prone to motivational effects.

The response vigour measures of *LFreq* and *LRel* confirmed that motivation changed in a quantifiable, consistent pattern between all animals to provide a readout of motivational level during a session. Next, we sought to determine whether an analysis of lick timing might allow us to gain a more nuanced insight into the behavioural strategy governing the “activating” licking rates.

### First-lick latencies reveal the balance of Impulsive and Instrumental influences on behaviour

While the licking frequency and the distribution of lick rates within specific trial sections reflect the “activational” aspects of motivation, the latency of an instrumental response can probe more deeply into the decision making process. Latencies have been used as indicators of behavioural impulsivity (Bari and Robbins 2013; Mayrhofer et al. 2013; Mayse et al. 2014) and the ability to inhibit inappropriate responses in decision-making tasks (Mayse et al. 2014).

We calculated first lick peri-stimulus-histograms (PSTHs) pooling latencies from all trial outcomes to identify the general trends of lick timing for a behaviour session. Figure 5a shows these group latencies for the monocular test session. We decomposed the response into two episodes: a primary lick response, defined by a sharp early peak immediately after stimulus onset, and a broad, multi-peaked secondary response which occurred at the end of the Buffer period, clustered around the 0.5s time-bin. The sharpness of the primary peak indicates a more stereotyped behaviour that is inflexible from trial to trial, typically seen in Pavlovian and overtrained habitual instrumental responses due to the availability of precise temporal information. To avoid habitual responses, animals were taken off daily training regimens immediately after attaining criterion and were kept on a reduced “reminder” regimen until testing. Comparisons with previous cohorts who formed habits after training for longer periods on easier stimuli confirmed the success of this strategy (data not shown). In the case of rodent lick-based tasks, licks can represent both an active indication of choice or the manifestation of an acquired ‘adjunctive’ consummatory behaviour, or Pavlovian response (Rescorla and Solomon 1967; Falk 1971; Schultz 2006; Dayan and Berridge 2014). Our use of the term Impulsive (which we could also refer to as Pavlovian within the scope of anticipatory and consummatory behaviours that are directionally aligned towards an appetitive goal), is intended to capture the broader relevance of these results to other experimental paradigms where choice and consumption responses are differentiated.

**Figure 5.**
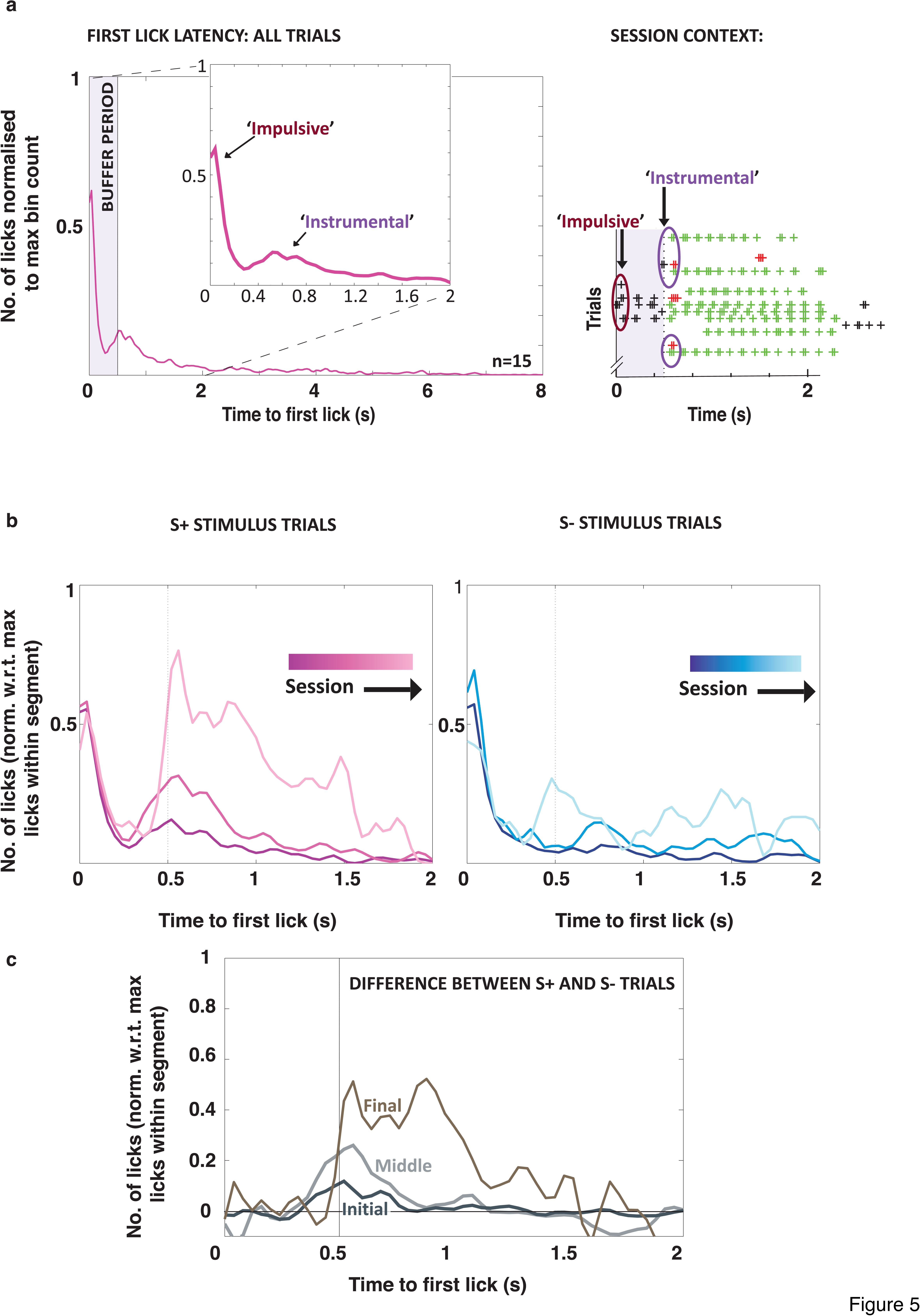
First-lick latencies reveal Instrumental and Pavlovian components of behaviour. ***a***. Left, peri-stimulus time histogram (PSTH) of first licks collapsed over all outcomes for the whole session for all mice (n=15). The distribution is bimodal with an immediate sharp peak (the Pavlovian component) following the trial onset, followed by a more dispersed secondary peak (the Instrumental component) at the end of the Buffer period (0.5s after trial onset). Inset: Enlarged view of the first-lick responses over the first 2s of the trials, providing the lick profile for just FA and Hit trials. Right, Part of a session report highlights some of the licks that contribute to the two prominent peaks seen in the lick profile. ***b***. The changes to first-lick profiles over the course of a session, calculated for the Initial, Middle and Final segments for S+ and S-trials, left and right respectively. (Initial = dark line, Final = pale line) Dashed arrows highlight the emergent changes to the post-Buffer lick profile towards the end of the session. The proportion of first licks occurring as part of an Instrumental response increases over the course of the session. ***c***. The differences between the lick profiles for S+ and S-trials for the Initial (dark grey), Middle (light grey) and Final (brown) segments obtained by subtracting the S-lick profiles from the S+ (raw values were used). *PSTH values were normalised to the time-bin with the highest number of first licks over the whole session (a) or for that particular session segment (b, c). All PSTHs were convolved with a Gaussian filter for smoothing, hence capping of highest peaks below 1 (see Methods for detail)*.

The spread in lick latency values seen during the secondary response on the other hand reflects more flexible decision-making. In this case, an animal’s Go/NoGo choice is based on the prior knowledge of the task contingencies in view of the trial-specific stimulus (S+ or S-) that is shown. We therefore describe the primary response as an “Impulsive” contribution to the behaviour and the secondary response as the “Instrumentally Driven” component. As the main features of the PSTHs were observed within the 0-2 sec time range (ie, during the Stimulus state), we focussed on comparing the latency profiles for the two outcomes where the animals made a ‘Go decision’ (Hits for S+ trials and FA for S-trials) and therefore necessarily made their first lick before 2 sec.

In order to relate the differences to our other performance indicators, we split the sessions into 3 equal segments as with the previous analyses (Figure 5). We extracted all Hit and FA trial latencies from within each segment and used a time range of between 0 and 2 seconds to construct PSTHs for the lick latencies. The lick latency PSTHs for both the Hit and FA trials changed considerably over the course of a session (Figure 5b). The difference between Hit and FA licks (Figure 5c) reveal that responses during these two trial types increasingly diverge in the secondary response (post buffer period), towards the end of a session. There is little difference between the primary response for both trial types, which might be expected given that licks during the buffer period had no consequence. This implies that the lick responses of the mice during the earliest part of the session show little difference irrespective of the stimulus presented (dark grey line in Figure 5c) and that discriminative licking patterns emerge later in the session (lighter grey lines, Figure 5c).

Peaks in the PSTH indicate the time at which it was most common for mice to register their initial lick, relative to the maximum response. The latency of the Impulsive response remained stable at 0.04 sec throughout the session. For the Instrumental component, latencies changed over the course of the session from 0.52 sec to 0.56 / 0.84 sec for Hit trials during the Initial, Middle and Final stages respectively. The differences in the timing of the two response components reinforces the suggestion that they are supported by different underlying processes, with the primary component showing more stereotyped, automated timing, while the secondary component remains prone to variation, suggesting a flexible decision making process. Interestingly, the latency of the peaks for the secondary component of the FA trials showed two widely separated modal peaks, at 0.48 sec and 1.44 sec. These groupings suggest two underlying error types, ‘Failure to Stop’, and ‘Failure to Wait’, which were previously reported by Mayse et al (2014). Such nuanced errors are indicative of a cost/benefit judgement of performing an action. The emergence of these lick latencies towards the end of the task reinforces the idea that mice start to rely on a behavioural strategy to solve the task in a goal directed manner.

We calculated the ratio between the peaks of the secondary and primary responses to quantify which grouping (Impulsive or Instrumental) contributed more to first-lick responses. To estimate the ratio we first quantified the licks within the Impulsive and Early Instrumental components by calculating the area under the curve for the lick PSTH between the times 0 - 0.2 sec and 0.4 - 0.6 sec respectively. The Impulsive component of the behaviour dominated the Initial segment for all mice, and this was reflected in the ratio of 2.88 for the peaks of the two response types (~3 Impulsive first licks for every 1 Instrumental first lick). By the Middle segment, the ratio shifted to 1.39 reflecting the shift to goal-directed behaviour. Nearer to the end of the session, Instrumental licking behaviour began to dominate (ratio=0.76). Over the course of the session, the Impulsive component gradually became less prominent, and the most frequently occurring first lick values belonged to the second, Instrumental grouping. Although this result might imply that the behavioural strategy is only optimised in the Final segment, an examination of the participation levels of the animals suggests otherwise. Namely, while the separation between Hit and FA values remains large and gives an appropriately high sensitivity value in the Final segment, the high proportion of Misses (71% (±0.05)) demonstrates that the mice were no longer well engaged in the task.

Overall, our analysis of the lick latency distribution revealed that as motivation changes, the balance of the behavioural processes contributing to decision-making shifts between impulsive and Instrumentally determined actions. Thus a highly motivated state is dominated by an Impulsive component, which obscures the optimal performance that Instrumental actions would otherwise result in. As a final test that high levels of motivation are in fact detrimental to performance on a discrimination task, we compared the performance of a mouse on consecutive days under satiation and deprivation.

### Deprivation may facilitate learning but prevents optimal performance once a task is learnt

Both the simulated results of the Actor-Critic “learning” (Figure 3b) and the instability of the ROC trajectories during learning (Figure 3c) suggested it was unlikely that the stereotypical poor performance of the mice during the Initial phase could be explained by learning. Furthermore, the Actor-Critic model predicted that performance should be stable over the course of a behaviour session if motivational state does not play a determining role (clustering of Hit/FA pairs in top panel of Figure 3b). To further test our hypothesis of the behaviour being motivation dependent, we compared the performance on consecutive days under a state of relative satiation on day 1 and normal deprivation on day 2. Figure 6 gives a typical example of these conditions for a single mouse to demonstrate the persistent vulnerability of Instrumental behaviour to “masking” by motivation irrespective of prior performance.

**Figure 6.**
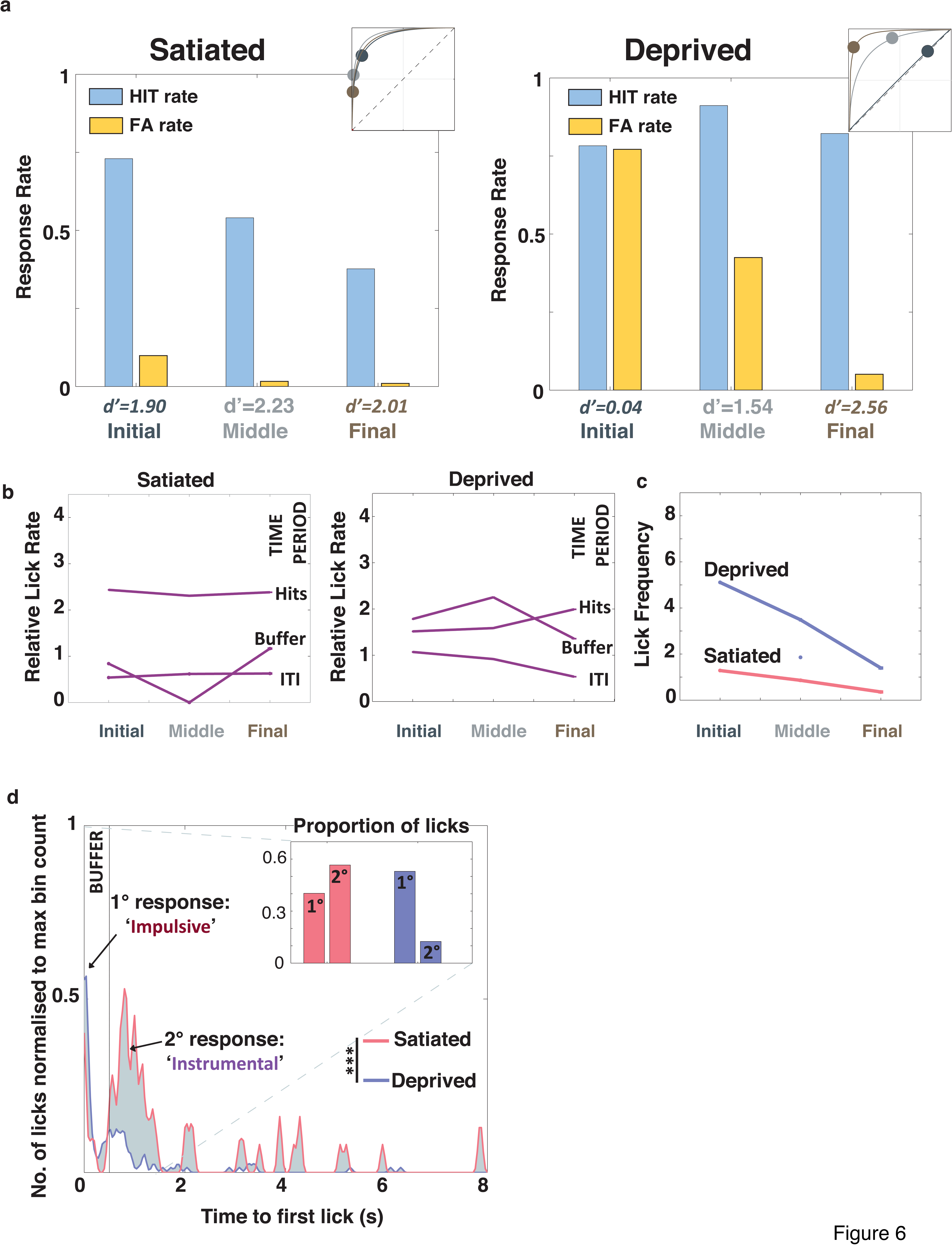
Satiated states result in better performance once a task has been learnt. ***a***. ROC plots for consecutive sessions by one mouse in the absence of (Satiated, 114 trials) and vulnerable to changes in motivation (Deprived, 168 trails). A brief free-water period prior to the Satiated session resulted in a stable instrumental performance throughout. Returning to the usual water restriction regimen for the Deprived test session on the following day reinstated the vulnerability of instrumental responding to motivational influence. ***b***. Relative lick rate (*LRel*) for 3 different time periods within trials are shown for this mouse: Hits, Buffer, ITI (see Figure 4b). Left, during the Satiated session, lick efficiency for Hits remained high throughout the session. Right, Buffer *LRel* remained during the Deprived state until the Final segment of the session. ***c***. The frequency of licks (*LFreq*) for the Deprived state followed the typical trajectory shown by the group, gradually decreasing over the course of the session while the Satiated *LFreq* showed little difference between the 3 session segments. ***d***. First-lick latencies (*LLat*) over all outcomes for the whole session. The majority of the first licks for the Satiated session occurred in the post-Buffer time period, suggesting the dominance of goal-directed responding. In contrast, the latency profile of the Deprived session showed a typical sharp peak immediately after trial onset and a much smaller secondary peak centered on 0.5s. (Satiated vs Deprived *LLat* profile, *** p<0.0001, Student’s t-test). 1°: primary, 2°: secondary peak. Inset: the difference between the Pavlovian and Instrumental components for both sessions.

Once criterion was reached, we established satiation by pre-feeding. Pre-feeding has previously been described as an effective short-term manipulation of motivation for reward incentive and responding (Salamone et al. 1991; Aberman and Salamone 1999). During the first session from the ‘Satiated’ state (Figure 6a, left) we found that the Instrumental component of the behaviour remained stable, with high *d*′ values throughout (*d*′=1.90-2.01). In contrast, when we returned to normal procedures for the session on the following day (Figure 6a, right), we confirmed that over-motivation affected even the best performing animals during the ‘Initial’ segment of the session. There was a negligible difference between the Hit and FA rates at the start of the session (Figure 6a, right: Initial), indicating that the animal was responding without any differentiation between the two stimuli (*d*′_Initial_=0.04, Hit_Initial_=0.78, FA_Initial_=0.77). Once the influence of motivation normalised, the animal returned to levels of response and performance indicators of a level that had been previously demonstrated *(d′*_Middle_*=* 1.54 *d*′_Final_=2.56). We also assessed the lick-based and response bias measures for the two sessions. As expected, the ‘Deprived’ session (Figure 6c) followed the same trajectories as described in the group results (Figure 4a). The session under relative satiation, in contrast, showed remarkable stability over the lick-based indicators *(LFreq* and *LRel)* indicating that the motivational influences described for the cohort were much reduced or completely absent. Specifically, the slope of the decrease followed by the *LFreq* over the 3 segments was minimal (Figure 6c), there was no crossover for *LRel*_Hit_ and *LRel*_BUFF_ (Figure 6b).

We also explored the relative contributions of the Impulsive and Instrumental components to the first-lick latencies for ‘Satiated’ and ‘Deprived’ sessions. The profiles of the PSTHs (for the whole session) were noticeably different (Student’s t-test: p<0.01) despite the relatively low number of trials contributing to the statistics. An examination of the general trends revealed that for ‘Satiated’ states, the majority of licks occurred in time bins after 0.5 sec and were likely to be goal-directed (main panel and inset, Figure 6d). This second response was multi-peaked with the majority of the maxima occurring between 0.7–1 sec. The ‘Deprived’ session, showed a considerably different pattern on the other hand (grey shading in Figure 6d highlights this difference between the two sessions). As with the ‘Satiated’ session, there was some evidence of a secondary response grouping close to 0.5 sec. These latencies correspond to the responses of the mouse that occurred in the Final segments towards the end of the session (data not shown). The primary response directly after trial onset, however, holds proportionally more of the latency values from the whole session. This grouping captures the Impulsive licking that the animals engage in when overmotivated. The inset of Figure 6d shows the absolute heights of the two response groupings. The ‘Satiated’ session had a dominant secondary response and a reduced Impulsive component. During ‘Deprived’ conditions, on the other hand, the contribution of Impulsive licking was proportionally much greater than the Instrumental.

## Discussion/Conclusion

We have shown that overmotivation can impair performance on a learnt visual discrimination task and, if performance is assessed by an average across a session, the true sensitivity of mice can be masked. The consistency of the changes to performance across subjects demonstrates that motivation affects behaviour in a stereotypical and predictable manner.

Our demonstration of the performance of a trained animal over consecutive days with ‘Satiated’ and ‘Deprived’ starting states respectively, shows that degradation in discrimination performance at the start of sessions is caused by overmotivation. Our results suggest that while deprivation may be necessary to drive initial learning during the training phase of a behavioural task, it can be detrimental to performance once a task is learnt. A detailed characterisation of changes to the distribution of first-lick latencies gave further insight into the cause of the variation to performance. The bimodal distribution of first-licks suggested that this variation was due to differences in the relative contribution of Pavlovian and Instrumental factors in states of high versus low motivation. At the start of a session, when animals are overmotivated, their actions are almost entirely driven by Pavlovian mechanisms, with only a small Instrumental component. In contrast, at the end of the session, the balance shifts, with most licks falling into the goal-directed grouping, but engagement with the task often at a relatively low level. Optimal performance is realised during the middle part of a behaviour session, where both Pavlovian and Instrumental responses are engaged to a high level, resulting in high participation and good discrimination. We conclude that the best performance on discrimination tasks requires a combination of these components.

By comparing the average *d*′ for the session with a segmented (within-session) analysis (Figure 2b) we were able to show that the former, which is the typical performance indicator used in behavioural neuroscience (Lee et al. 2012; Glickfeld et al. 2013), misrepresents the true abilities of mice on a GO/NOGO task. Averaging the *d’* over the session underestimates the mouse’s discrimination capability, due primarily to reduced sensitivity in the initial part of a behaviour session. Instead, we recommend fitting *d’* to data collected across the session, excluding the early (over-motivated) trials, by tracking the animal’s progress across the ROC throughout the session in a sliding or shifting window. This builds on previous suggestions of rank-biserial correlation between trial number and responses (Stüttgen and Schwarz 2008) and modelling by Wichmann and colleagues (Wichmann and Hill 2001; Fründ et al. 2011) that suggested statistical methods to overcome non-stationarity.

A comparison of 2 different modifications of the popular Actor-Critic model (Dayan and Balleine 2002; Niv et al. 2007; Dayan and Niv 2008) demonstrated that the performance curves observed in our behavioural data were most closely replicated by introducing a motivation variable into the framework. This further supports the idea that the “masking” of high discrimination ability is caused by short-term motivational influences, rather than the persistence of a dynamic learning phase throughout the testing period. Guided by the previous work of Salamone & Correa (2002, 2012) and Niv et al (2007), we quantified the activational aspect to motivation with the licking frequency, showing a linear decrease over the course of the session. We also showed that licking became differentially more focused towards the rewarding times within the trial (*LRel*_Hits_ vs *LRel*_Buff_ comparison, as has previously been suggested with a different behavioural paradigm with rodents (Komiyama et al. 2010). We consider it likely that a Pavlovian response is enhanced by the display of the visual stimulus at the start of trials, as it has previously been suggested that Pavlovian responses can undergo a generalisation effect that dissociates them from the specific details of a stimulus (Dayan and Berridge 2014). Therefore, licking under Pavlovian influence should be disproportionately increased during the Buffer for all stimuli in the early stages of a session. Towards the end of a session, actions (licks) are deployed in a more strategic manner, supporting our claim that the balance of behavioural strategy shifts towards goal-directed as the trial progresses.

As such tasks become more widespread, the behavioural analyses described here may be a useful resource for neurophysiologists investigating the neurobiological basis of behaviour. Our results may help with task design and appropriate analysis constraints. A growing number of groups are using Go/NoGo paradigms with SDT performance measures. Such studies use neurometric readout and psychophysical performance to interpret perceptual thresholds and sensory coding (Sugrue et al. 2005; Stüttgen and Schwarz 2008; Nienborg et al. 2012; Carandini and Churchland 2013). Our behavioural results show that the ongoing balance of Pavlovian and Instrumental factors is likely to shape the extent to which an animal relies on sensory information in order to guide its behaviour (Stüttgen et al. 2011). Licking behaviour is but one example of a commonly employed instrumental action that suffers from the difficulty of separation between Pavlovian and instrumental components, showing the wider relevance of our results. Approach behaviour towards appetitive reinforcers can interfere with the interpretation of paradigms that use spatially separated levers and/or maze-choice arms in operant chambers for freely moving animals (Smith et al. 2012; Goltstein et al. 2013). A recent study using human subjects has also described similar Pavlovian to instrumental imbalance in decision-making (Chumbley et al. 2014). We hope that these results can guide others towards the accurate classification of behavioural components in order to be confident in their interpretations of neurometric readout. By using quantifiable measures of the “activating” aspects of motivation, researchers can set thresholds and decide on reproducible and consistent criteria, that best suit the question they are investigating to systematically compare behaviour between groups or within individual animals.

Future work might expand on the measures presented by our group to consider alternative readouts of motivational state. This could include further analysis of session data, for instance by the use of breakpoint analysis. From the perspective of behaviour as a whole-body process, this might also involve monitoring physiological variables such as pupil diameter, muscle tension and heart rate, in order to optimise the behavioural regime for each subject and session.

## Methods

### Subjects

All experiments used female C57BL/6 (Harlan or Charles River Laboratories) mice that were 4-6 weeks of age at the time of headplate surgery (n=15). Animals were group housed on a reversed 12 hour light-dark cycle and experiments were carried out in the dark phase. Mice were weighed and handled daily throughout water restriction. Food was available *ad libitum* and animals received daily water supplementation either during behavioural training or in their home cage on rest days to maintain weight above 85% of the starting value. All procedures were carried out in accordance with UK Home Office Animal Care Guidelines, and approved by the Imperial College Animal Welfare and Ethical Review Body under Project License 70/7355.

### Headplate implantation

Prior to water restriction and behavioural training all animals underwent a headplate implantation surgery. All surgeries were carried out using aseptic technique under isoflurane anaesthesia. Throughout surgery, body temperature was monitored with a rectal probe. Immediately after induction of anaesthesia, 0.1ml of carprofen (1mg/ml, Rimadyl©) was injected subcutaneously. After exposing the skull, a drop of 5% hydrogen peroxide was used to clean the skull surface. Custom designed metal headplates were attached using Histoacryl^®^ (TissueSeal LLC, TS1050071FP), leaving access to a 5mm diameter site centered on monocular V1 (2.5-3.2mm lateral to lambda) (Lee et al. 2012; Glickfeld et al. 2013). The craniotomy site was filled with silicon elastomer (Kwiksil ™, World Precision Instruments, Inc) and covered with a layer of coloured nail varnish to discourage the group-housed animals from interfering with the site prior to electrophysiology experiments. A small surgical screw (Precision Technology Supplies, M1.2x2.4) was implanted contralaterally to act as a ground. The headplate attachment was reinforced with dental acrylic and the cut edges of the skin were re-attached using Histoacryl. Throughout the surgery, the eyes were covered with Vaseline or eye ointment to prevent drying. Following the procedure, animals were transferred to a heated (~35°C) incubation chamber, where they were allowed to recover to normal activity with ad libitum water and food access. The water restriction protocol was started 1-2 days following surgery.

### Behavioural apparatus

We used a 800mm (i.d.) polystyrene toroid (Ecclestone & Hart Ltd) to create an immersive visual environment based on the designs of previous virtual-reality environments (Hölscher et al. 2005; Harvey et al. 2009). All programming of training and behavioural paradigms used LabVIEW System Design Suite coupled to a National Instruments Data Acquisition Board (NI-DAQ). Each sub-division of the task (task state) was matched to a visual background projected onto the interior of the dome. Visual backgrounds, including the equiluminant GO and NOGO stimuli, were generated using an OpenGL framework with C++ syntax. The resulting texture was mapped to a polygon on the computer screen using specified coordinates of a grid-like mesh to the dome interior (Tomazzo Muzzu, in preparation). A digital projector (ViewSonic PJD 6533w, 120Hz refresh rate, output brightness 3500 Lumens) relayed the dome-adjusted image onto a rectangular mirror mounted at 35°, which reflected onto a 180° spherical mirror (Viso MS180 Indoor Dome Mirror) to ensure the image was projected onto the appropriate coordinates of the dome interior (Figure1). Water rewards were delivered through a blunted 19G syringe needle attached to 0.5mm bore silicone tubing (RS Components, Stock No.667-8438). Water delivery was controlled by a TTL pulse to a single channel peristaltic pump (Campden Instruments, Product No.80204-0.5). Licks were detected with a photomicrosensor (Omron Electronics, EE-SX4070) sampled at 50Hz. All interruptions to the beam emitted by the sensor were counted as licks. Airpuff delivery was controlled by a 2-2 way solenoid valve (Shako Company, PU220AR-01, ⅛ inch) connected to an air pressure regulator, allowing manual adjustment of air pressure (10-15psi). Airpuffs were delivered via a 1mm copper piping angled towards the mouse from the side opposite to the monocular projection.

### Training paradigm

Time spent in each phase of training was allocated on an individual basis governed by experimenter experience and pre-determined criteria. Throughout training and testing, the number of trials per session typically ranged between 100-300 trials. Length of session was determined by mouse participation levels; sessions were terminated after ~10-20 non-participatory trials. These non-participatory periods were not included in the analysis. ***Pre-training phase***: Mice were gradually habituated to head fixation over several days, starting with 5 minute sessions. ***Autoshaping:*** The S+ stimulus (reward-coupled) was a leftward drifting squarewave vertical grating (Temporal Frequency (TF): ~3.3 Hz). Only S+ was shown at this stage, alternated with blank (black) screen every 15 trials. For the first session, water reward was automatically delivered following the presentation of S+, before transitioning to the instrumental contingency where reward state was lick-triggered by the animal. Licking during the blank screen presentation was punished with an airpuff (0.3-0.5 sec) to the upper body to prevent compulsive over-licking. Mice progressed to the next stage when they displayed higher lick efficiency by targeting their licks to water rewarded time periods of the trial. ***Full task (Full-field):*** S+ and S-were alternated in a pseudorandom pattern determined by a random number generator in LabVIEW with a rule preventing the presentation of the same stimulus more than three times in a row. The S-stimulus (airpuff-coupled) was a downward drifting horizontal grating (TF: ~0.5 Hz). Progress during full-task training was evaluated using Signal Detection Theory (SDT) measures. When high discrimination sensitivity was observed in three of five consecutive sessions, mice entered the final phase of training where a monocular projection region was used (outer ~60-135° of the right visual field). Despite similarity of full-field and monocular stimuli (same SF and TF), performance level of the animals typically dropped at this stage of training. Animals took much longer to reach criterion performance levels, in some cases requiring adjustment of stimuli parameters (TF) to make them more easily discernable. Criterion performance was evaluated using a combination of mean/optimal <u>d’</u> values and observing the profile of Correct Rejection rates over session. When high sensitivity (as indicated by d’ values >1.0) was observed in three of four sessions, the mice were considered well-trained. After reaching criterion animals were taken out of the daily training regime to prevent overtraining to habitual responding. Thereafter, animals were kept on a regular “reminder” regime limiting sessions to one every 2/3 days, which enabled us to ensure a high level of performance on the task. These criteria were established in accordance with current behavioural literature (Komiyama et al. 2010; Smith et al. 2012; Horner et al. 2013; Rivalan et al. 2013; Izquierdo et al. 2013).

***“Satiated” state:*** Satiation was established by pre-feeding, which consisted of free water access for 1-2 minutes before the behaviour session.

***Timings and State Flow of Task:*** Each trial started with a Stimulus State (2 sec total), which consisted of an initial 0.5 sec “Buffer period” and a 1.5 sec “Response Window”. During the Buffer, licks made by the animal did not count towards a decision/state change. The Buffer period was used within the trial structure to allow mice enough time to attend to the stimulus in view of the rapid succession of trials, and for mice to have time to bring licking under control, as guided by previous behavioural task literature (Lee et al. 2012; Pinto et al. 2013; Bracey et al. 2013). For every single mouse, FA trials had more licks in the Buffer period than Hit trials (see Supplementary Figure 1). S+ trials qualified as Hits when the animal licked within the Response Window of the stimulus presentation, and Misses otherwise. During the Reward State (1.75s), S+ continued to be projected onto the screen while a 1-4 μl water reward was delivered to the mouse. S-trials were designated as Correct Rejections (CR) if no licks occurred during the Response Window and False Alarms (FA) otherwise. Punishment for FAs consisted of a mild airpuff (0.5 sec) to the upper body and a 5 sec timeout period while a black screen was projected onto the dome. Any licks during the timeout period triggered additional brief (10 ms) airpuffs. Misses and CRs were not negatively or positively reinforced. A variable length (3-6 sec) Inter-trial Interval (ITI) State immediately followed the Reward, Punishment or Miss/CR States.

***Data analysis:*** All analyses were carried out in MATLAB using scripts developed in house and built-in functions. Over a given session, a sliding window over 30 trials was used to calculate Hit and FA rates (step size: 1 trial). Hit rate was calculated as (Hits/ total S+ trials) and FA rate (FA/total S-trials). We clipped values of Hit and FA rates that were >0.99 or <0.01 (Miller 1996). Performance was evaluated using the SDT sensitivity measure (*d’*) from the inverse normal cumulative distribution of the Hit and FA rates: *d’ = z(Hit) - z(FA)*. We split sessions into 3 segments (Initial, Middle, Final) and projected Hit/FA pairs into Receiver Operator Characteristic (ROC) space. Statistical tests were Student’s t-test for parametric pairwise and unpaired comparisons and repeated measure ANOVA with post-hoc comparisons; error bars show s.e.m.

***Lick latency peri-stimulus-time-histogram (PSTHs):*** We calculated histograms (time bins = 0.04 sec) of first licks per trial for the group. All PSTHs were convolved with a Gaussian filter for smoothing and normalized to the maximum bin count for the session (whole session PSTH) or for the segment (trial outcome PSTHs) to determine the relative likelihood of licks occurring within particular time bins. We classified the bimodal distribution of the lick latencies, into primary (1°) “Pavlovian” (0 - 0.2 sec) and secondary (2°) “Instrumentally Driven” responses (0.4 sec+).

### Reinforcement Learning Modelling

To model behavioural responses, we used the classic Q learning model (Watkins 1989, Sutton and Barto 1998). In this model, the agent takes probabilistic actions in certain states, given a set of real values called Q-values. Our version of the model uses two actions (L, NL) and two states (GO, NOGO). The states correspond to two different stimuli, S+ and S-, for Go and NoGo respectively, and the two actions are L (Lick) and NL (No Lick). In total there are two Q-values in each time step, each corresponding to a given state (Q_1,t_:S+ at time t and Q_2,t_:S-at time t). The higher the Q-value the more likely the agent will take the lick action. Conversely the lower the Q-value the less likely the agent will take the no-lick action. To model mouse decision making, we used the non-deterministic Softmax decision rule,

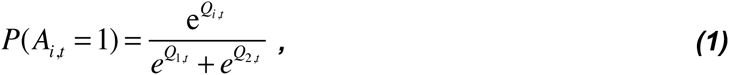

where p(*A*_*i,t*_=1) is the probability of taking the lick action given that the mouse is in the *i*-th state at time *t*, *Q*_*i,t*_ is the value of licking in the *i*-th state in the *t*-th time bin, and T is a “temperature” parameter that was fixed to 1 in all our simulations. A session corresponds to N=150 trials where the first action is randomly assigned to 0 or 1 with a probability of p=0.5. We used R=15 repetitions of this protocol and computed the hit and false alarm rate each time. This choice of parameters reflected the experimental protocol.

We compared two different models, which we have denoted AC and MAC. The AC model is the classic actor-critic Q-learner, which learns at the single session level, with the Q-values being modified as

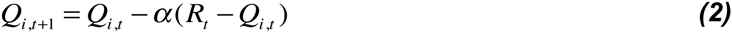

where *Q*_*t*_ is the value at a given time *t*, α is the learning rate, and *R*_*t*_ the reward provided to the agent at time *t*. This updated rule guarantees that the Q values converge towards the mean reward when the agent takes the action in this state. For AC model, the reward is 1 when the mouse licks during a GO stimulus or when it refrains from licking during a NOGO stimulus, otherwise the reward is −1. Figure 3a shows results from the AC model where Q-values are initialized to 1 under the assumption that the mice have learnt the task contingencies during discrimination training.

The MAC (Motivated Actor-Critic) model is similar to AC in all respects, except that we have introduced a real valued variable *M* which accounts for the motivation or thirstiness of the agent. *M* diminishes by a fixed step *m* each time the agent received a reward. The Q-values we used are now biased by a motivation variable *M*,

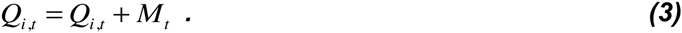

M is modified if the mouse receives a liquid reward during a GO stimulus:

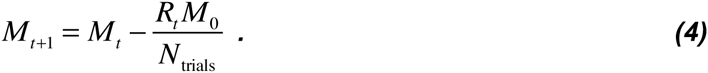

The code used to generate Figure 3 is available as a module (motivBCS2015) that can be downloaded from our website (link to be provided). All the parameters used in our model are presented in Table 2.

**Table 2.** Model Parameters.

| Parameter | Value |
| --- | --- |
| $p$ | 0.5 |
| $N$ | 150 |
| $R$ | 15 |
| $T$ | 1 |
| $Q_{\text{init}}$ | 0 |
| $\alpha$ | 0.1 |
| $M_0$ | 1 |
| $\alpha_m$ | 0.06 |

## Author contribution statement

A.B. performed the experimental work described, R.D.C. performed the modeling work described, A.B. prepared the figures, and A.B., R.D.C. and S.R.S. wrote the manuscript. All authors reviewed the manuscript.

## SUPPLEMENTARY INFORMATION

**Supplementary Figure 1.**
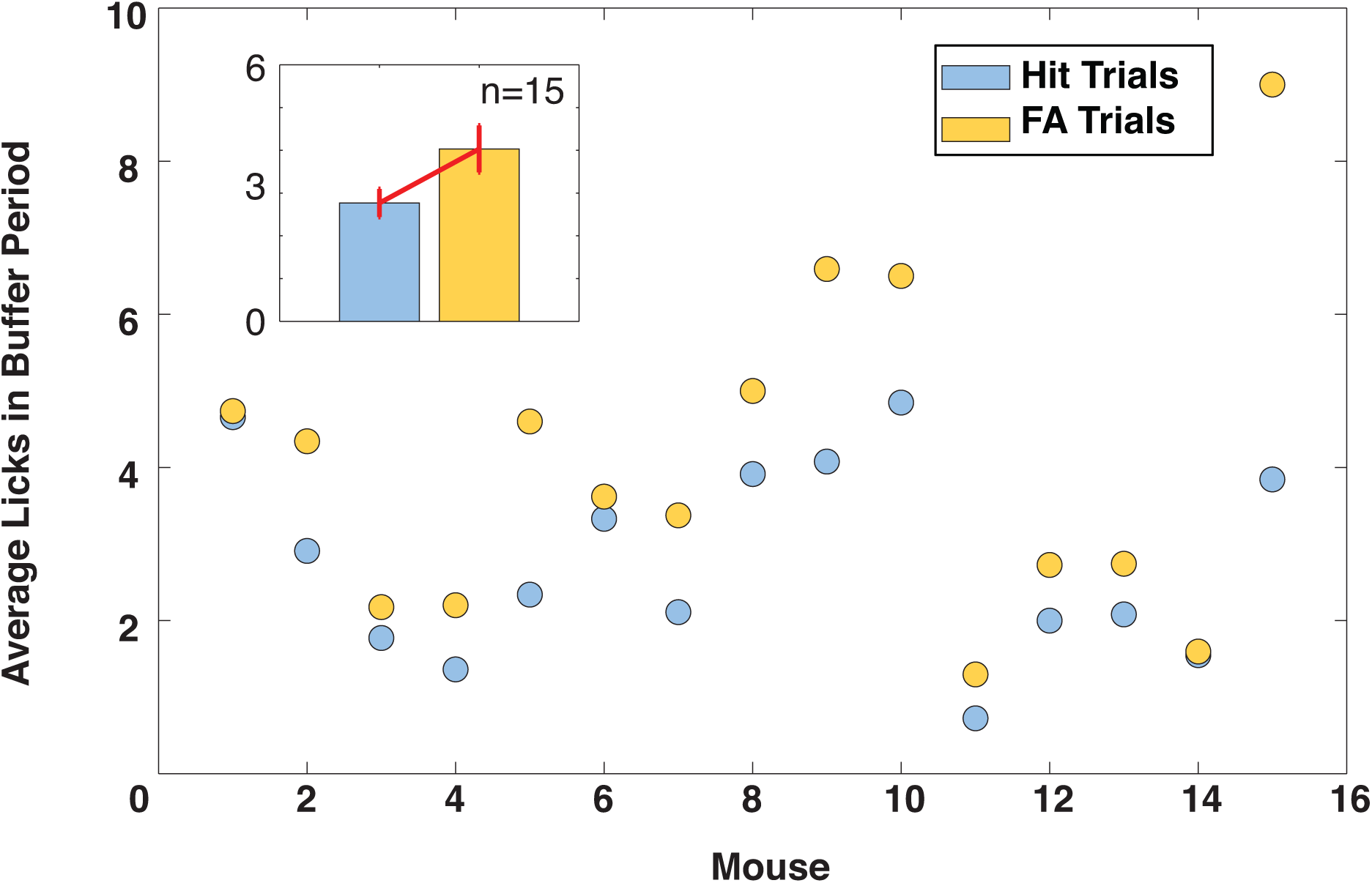
More licks occur in the Buffer period of FA trials than Hit trials. Main panel: The figure shows the average number of licks in the Buffer periods for Hit (blue) and FA (yellow) trials for each mouse. Inset: The average number of licks in the Buffer period for Hit and FA trials for the cohort (mean +/- s.e.m., n=15).

